# The folding space of protein *β*2-microglobulin is modulated by a single disulfide bridge

**DOI:** 10.1101/2021.05.19.444822

**Authors:** Jules Morand, Ana Nunes, Patrícia F.N. Faísca

## Abstract

Protein beta-2-microglobulin (*β*2*m*) is classically considered the causative agent of dialysis related amyloidosis (DRA), a conformational disorder that affects patients undergoing long-term hemodialysis. Together with the wild type form, the Δ*N*6 structural variant, and the D76N mutant, have been extensively used as model systems of *β*2*m* aggregation. In all of them, the native structure is stabilized by a disulfide bridge between the sulphur atoms of the cysteine residues 25 (at B strand) and 80 (at F strand), which has been considered fundamental in *β*2*m* fibrillogenesis. Here, we use extensive Discrete Molecular Dynamics simulations of a full atomistic structurebased model to explore the role of this disulfide bridge as a modulator of the folding space of *β*2*m*. In particular, by considering different models for the disulfide bridge, we explore the thermodynamics of the folding transition, and the formation of intermediate states that may have the potential to trigger the aggregation cascade. Our results show that the dissulfide bridge affects folding transition and folding thermodynamics of the considered model systems, although to different extents. In particular, when the interaction between the sulphur atoms is stabilized relative to the other intramolecular interactions, or even locked (i.e. permanently established), the WT form populates an intermediate state featuring a well preserved core, and two unstructured termini, which was previously detected only for the D76N mutant. The formation of this intermediate state may have important implications in our understanding of *β*2*m* fibrillogenesis.

## 1. Introduction

Human *β*2 microgobulin (*β*2m) is a small (*N*=99 residues) protein with a beta-sandwich native fold. The wild type (WT) form together with a structural variant termed ΔN6 (lacking the first six N-terminal residues) (Fig. 1), are associated with the onset of dialysis related amyloidosis (DRA), a condition that affects the bones and cartilages of individuals with kidney failure undergoing long-term (> 10 years) hemodialysis. On the other hand, the D76N mutant is the culprit of a systemic amyloidosis affecting visceral organs (liver, kidneys etc) [1]. The three model systems (WT, ΔN6, D76N) contain two important structural features that have been implicated in the onset of amyloidogenesis: A disulfide bridge (or SS-bond) between the sulphur atoms of two cysteine residues that contributes to stabilise the native structure (Figure 1), and a thermodynamically unfavourable, cis prolyl-peptide at position 32. Interestingly, the difference in thermodynamic stability between the two isomers is attenuated in the D76N mutant, which may facilitate the transition between the two forms [2].

**Figure 1.**
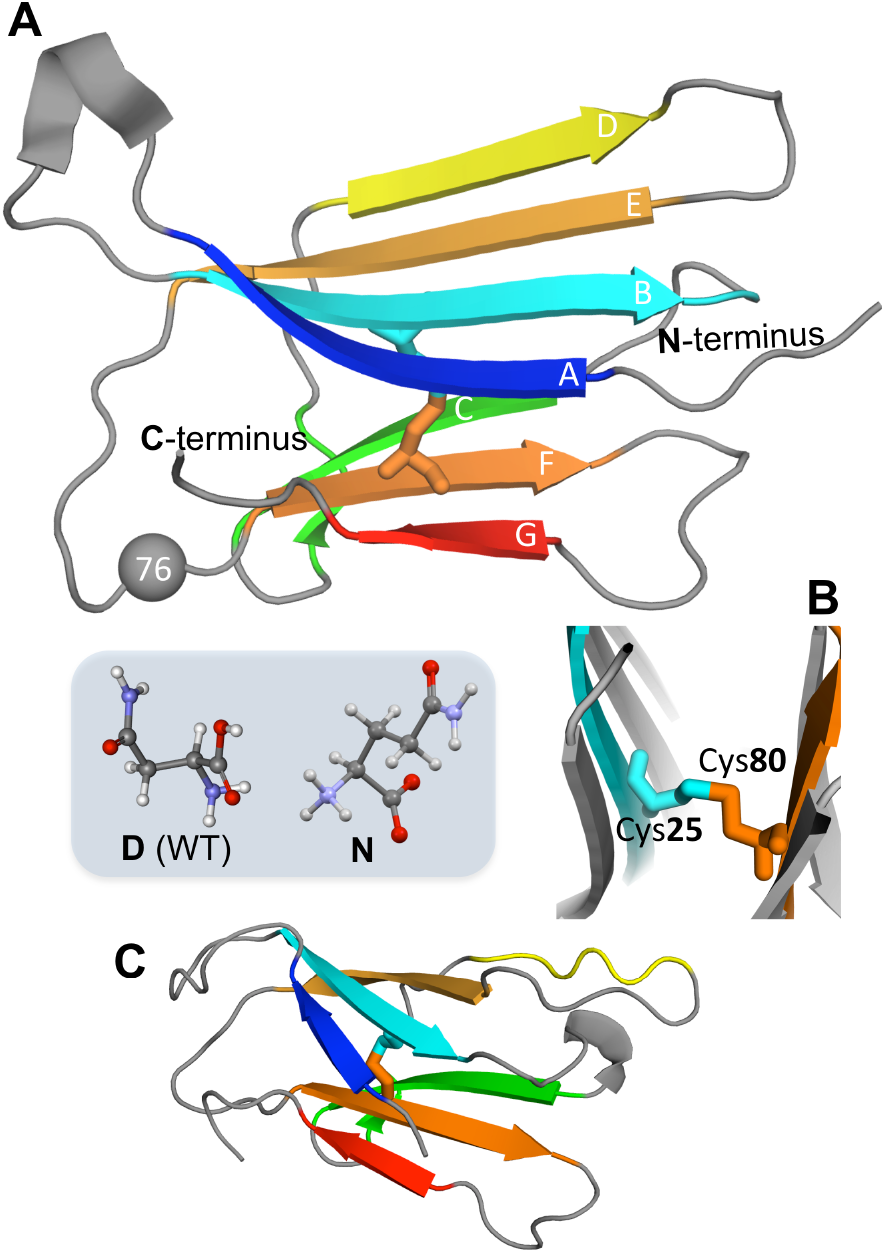
Protein *β*2-microglobulin. Three-dimensional native structure *β*2m highlighting the seven beta-strands, organised in two beta-sheets (comprising strands A-B-E-D, and C-F-G, respectively) (PDB ID:2YXF), and position 76, which is mutated to Asn in the D76N mutant (PDB ID: 4FXL) (A). The SS-bond between the sulphur atoms of residues 25 and 80 that links strands B and F (B). Native structure of the ΔN6 structural variant lacking the N-terminal hexapeptide (PDB ID: 2XKU) (C).

In principle, a cure for amyloid disease requires solving the aggregation mechanism. The latter is a remarkably complex process, which involves the formation of molecular structures of varying size and timescales, being highly dependent on environmental conditions. To solve the aggregation mechanism, one should be able to determine the size, distribution and structures of the oligomeric assemblies, filaments, protofibrils and fibrils that make up the amyloid pathway, as well as the rate constants associated with every transition [3]. Ideally, it should be possible to pharmacologically block the process as early as possible, but the transient lifetimes, and complex dynamic equilibrium according to which aggregation-prone monomeric species and early-formed oligomers coexist with each other, greatly challenges their structural characterisation with atomic resolution [4]. To complicate matters further, the WT form of *β*2*m* does not aggregate *in vitro* in physiological conditions without additives (reviewed in [5]).

Despite these challenges, progress has been made to establish the initial steps of *β*2m aggregation via experiments *in vitro* as well as through molecular simulations (reviewed in [6]). At the monomeric level, an important player of the aggregation mechanism has been identified and characterised. This folding intermediate is overall nativelike but features the prolyl-peptide bond at position 32 in a non-native trans conformation. For this reason it has been named I_*T*_. Experimental studies reported an equilibrium population of 5%-15% [7, 8] for the WT form, between 5%-25% for the D76N mutant [9, 10], and 90% for the ΔN6 variant [11]; the latter is therefore considered a structural mimic of I_*T*_.

The exact mechanism according to which I_*T*_ drives aggregation of the WT form is a matter of debate [5], and in the case of the D76N mutant conditions specific of the *in vivo* environment (e.g. shear forces) have been invoked to explain its role as trigger of the aggregation cascade [10]. The intermediate state I_*T*_ populated by D76N represents a highly dynamic structure [12], and conformational excursions to conformers with an higher aggregation potential are likely to occur. In line with this observation we recently proposed the existence of an intermediate, which we found to be exclusively populated by the D76N mutant, featuring a well preserved core, and two unstructured termini that expose highly aggregation prone beta strands [13, 14, 6]. The detachment of the terminal regions is consistent with the fact that the D76N mutation breaks a large network of electrostatic interactions distributed over a large number of residues including the C- and the N-termini. Our structural prediction, which is based on results from discrete molecular dynamics (DMD) simulations of a structured-based Gō model [15, 16, 17], is in line with results based on ssNMR according to which D76N sparsely populates a highly dynamic conformation that exposes very aggregation prone regions as a result of a loss of *β*-structure at the N- and C-terminal strands [18]. These results highlight the importance of structure-based models in predicting nativelike conformational states [19].

The role of disulfide bridges in protein folding and stability has long been an active topic of research [20]. A recent simulation study based on a coarse-grained model of protein bovine pancreatic trypsin inhibitor (BPTI), which has three disulfide bonds in its native state, showed that – at least for this particular protein – it is the folding process that directs disulfide bond formation, and not the other way round [21]. In the case of *β*2m, the role of cysteine bond between residues 25 and 80 in aggregation has also been investigated and several experimental studies indicate that the SS-bond is essential to observer amyloid fibril formation [22, 23, 24]. Mechanistic insights from DMD simulations also support the view that the SS-bond has a critical role in amyloidegenesis by showing that the morphology of the dimers formed when the cysteine bond is established is compatible with further oligomerization through parallel stacking that eventually leads to amyloid fibrils, while a mechanism based on domainswapping is observed in the absence of the cysteine bond, which should result into short, curved fibrils [25].

Here we go back one step and use DMD simulations to explore the role of the SS-bond at the level of the monomeric (i.e. folding) space of the three model systems. More precisely, we are interested in determining how this bond affects the folding transition, and the free energy landscape of *β*2m. We find that for the three investigated model systems (WT, ΔN6 and D76N) an increase in the strength of the SS-bond leads to higher thermal stability of the native state, and causes the folding transition to become less cooperative, specially for the WT form. We found that the SS-bond has the ability to modulate the free energy landscape by enhancing the population of intermediate states with potential to trigger the amyloid pathway. In the case of the WT form this effect translates into the population of the intermediate state *I*_2_, which had been previously been detected only for the D76N mutant. The results of the present study thus reinforce the importance of the SS-bond in *β*2m aggregation.

## 2. Model and Methods

### 2.1. Full atomistic protein representation and Go potential

We consider a representation of the protein where all non-hydrogen atoms are represented by hard spheres of unit mass. The size of each atom if defined by its van der Waals (vdW) radius [26] scaled by factor *α* < 1. To model protein energetics we consider i) excluded volume interactions, ii) bonded interactions, and iii) non-bonded interactions, which are all modeled by discontinuous, piecewise constant interaction potentials. To take into account excluded volume interactions, all atom-atom interactions at distances smaller than the sum of the hard-sphere radii, i.e. *r_ij_* < *α*(*r*_0*i*_ + *r*_0*j*_), where *r*_0*i*_ is the vdW radius of atom *i* and *r*_0*j*_ is the vdW radius of atom *j*, are strictly forbidden. Bonded interactions (i.e. covalent and covalent-like bonds) between adjacent atoms *i* and *j* are modelled by a infinitely narrow high potential well

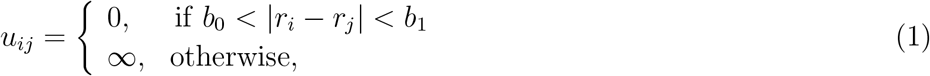

with *b*_0_ = 0.9 and *b*_1_ = 1.1 such that the average covalent bond is equal to 1Å.

Non-bonded (or contact) interactions are represented by a square-well potential whose depth is given by Gō energetics [27]. Thus, if atoms *i* and *j* are located in residues which are separated by more than two units of backbone distance the interaction parameter between them, *ϵ_ij_*, is given by

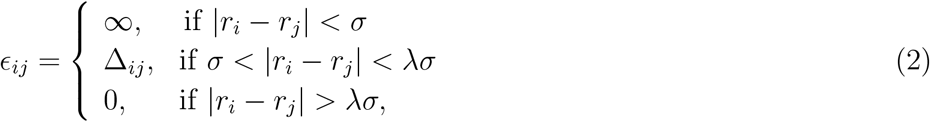

where *σ* = *α*(*r*_0*i*_ + *r*_0*j*_), λ a scaling factor controlling the range of attractive interactions, and Δ_*ij*_ = —1 if *i* and *j* are in contact in the native conformation and equals 0 otherwise. We set *α* = 0.80 and λ = 1.6 to have a well behaved folding transition. With this choice of parameters the cut-off distance is (for methyl carbons) 4.7Å. The total energy of a conformation is given by

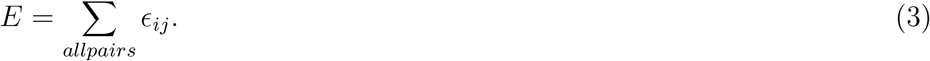

### 2.2. The disulfide bridge

The formation/rupture of a SS-bond is a process that involves the oxidation/reduction of the thiol groups belonging to cysteine residues. Thus, in principle, simulating the formation and rupture of an SS-bond in the context of a coarse-grained model is not necessarily straightforward, and several criteria (e.g. proximity and correct orientation of the S atoms, access of thiols to oxidizing agents) must be adopted to accurately capture this process [21]. Here, to account for the SS-bond between the sulphur atoms of residue Cys25 (located on strand B) and residue Cys80 (located on strand F), we consider three simple models: In the first, which we term *native interaction* model, the SS-bond is considered a non-bonded interaction as described by eq.(2). This means that the interaction forms and breaks during the course of a folding simulation, and contributes as much as any native interaction (i.e. *ϵ_ij_* = —1 energy units) to the total energy of a conformation. Assuming that a typical intramolecular hydrogen bond has an associated interaction strength of 5Kcal/*mol*, and setting the potential energy for a native h-bond to unit energy, the interaction strength of a SS-bond is obtained by normalising its typical interaction strength (60Kcal/*mol*) to the interaction strength of the h-bond. Thus, in the second model we consider, which we term *intermediate interaction* model, the SS-bond is taken as a stronger non-bonded interaction by increasing the depth of the potential well 12 times, so that the SS-bond contributes with *ϵ_ij_* = —12 energy units to the total energy of a conformation. Finally, in the third model, termed *locked interaction* model, the SS-bond is taken as a covalent bond, such that the two sulphur atoms interact via the narrow infinitely high potential well described above (eq.(1)). This implies that the bond does not break in the course of a folding simulation that starts from an unfolded conformation with the SS-bond established. Despite their simplicity, the adopted models should be able to qualitatively capture the effects of the SS-bond on the folding process of *β*2m.

### 2.3. Conformational sampling and data analysis

To sample the conformational space we use discrete molecular dynamics (DMD) simulations [28]. Folding progress is evaluated by computing several reaction coordinates such as the energy (*E*), fraction of native contacts (*Q*), root-mean-square deviation (*RMSD*) to the native structure, and radius of gyration (*R_g_*). Simulations start from unfolded conformations (*R_g_* > 30Å). In order to properly sample the conformational space at different temperatures, and observe multiple folding-unfolding transitions we use RE-DMD by combining the DMD engine with a replica-exchange (RE) Monte Carlo method [29]. Data for thermodynamic analysis is collected by sampling uncorrelated states from RE-DMD trajectories consisting of up to 3 × 10^10^ events after equilibration. The details of RE-DMD simulations can be found elsewhere [15, 19, 16, 17]. The heat capacity, *C_V_* = (〈*E*^2^〉 – 〈*E*^2^〉)/*T*^2^, was evaluated as a function of temperature, and the melting temperature, *T_m_*, was determined as the temperature at which *C_V_* reached its maximum value. Data from RE-DMD is analysed with the weighted histogram analysis method (WHAM) [30]. In particular, we use the WHAM method to compute free energy profiles (i.e. the free energy projected along *E*) and free energy surfaces (i.e. the free energy projected along *E, R_g_* or *RMSD* to native structure). We evaluate the transition temperature *T_f_* as the temperature at which the native and unfolded states are equally populated at equilibrium. We conduct long (up to 8 × 10^10^ events) DMD simulations at fixed temperature to get ensembles of relevant conformational states observed in the free energy profiles and surfaces. To determine the relevant conformational classes present in an ensemble of conformations we used the hierarchical clustering algorithm *jclust* available in the MMTSB tool set [31]. From each identified cluster we extracted the conformation that lies closest to the cluster’s centroid.

## 3. Results

### 3.1. Model systems

In the present study we use the high-resolution crystal structure of Goto and coworkers [32] as a model system for the WT form (PDB ID: 2YXF), and that of Bellotti and coworkers [12] as a model system for the D76N mutant (PDB ID: 4FXL). In the case of the structural variant ΔN6, we use the NMR-determined structure determined by Radford and co-workers (PDB ID: 2XKU)[33, 34]. The native contact maps are represented in Figure 2. The radius of gyration of the native structure are 14.36 A(WT form), 14.25 A(D76N mutant) and 13.85Å(ΔN6). It should be mentioned that the high-resolution crystal structures of D76N and WT form are very similar to each other, with a root mean square difference of 0.59Å[1].

**Figure 2.**
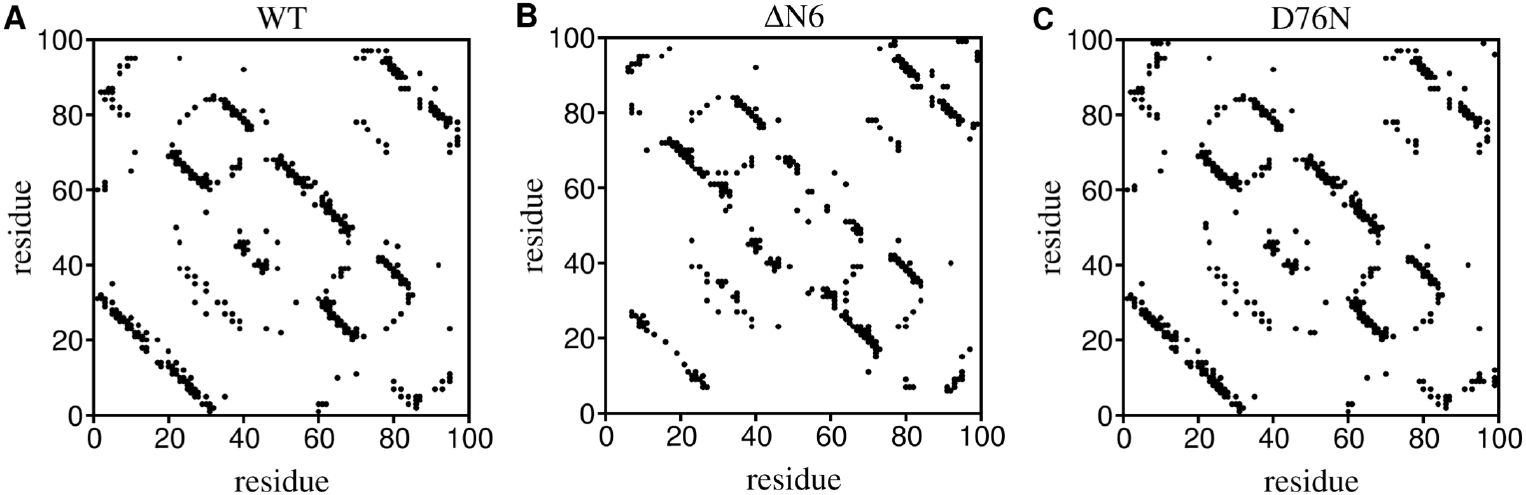
Contact maps. Two-dimensional *N* × *N* matrices representing the contacts that exist in the native structure of the WT form (A), D76N mutant (B) and ΔN6 structural variant (C). Within the context of the adopted Gō potential there are 1136 native contacts in the WT form, 1311 native contacts in D76N, and 899 native contacts in ΔN6, which translate into a native energy of *E_WT_* = —1136, *E*_*D*76*N*_ = —1311 and *E*_Δ*N*6_ = —899 energy units.

### 3.2. Folding transition

We started by evaluating the role of the SS-bond on thermal stability, as measured by the melting temperature *T_m_* (Fig. 3). Our results indicate that as the strength of the SS-bond increases, *T_m_* typically increases, in consonance with a higher thermal stability. The increase in thermal stability when the SS-bond is locked is particularly striking in the case of the ΔN6, with *T_m_* increasing by 11.3% with regard to the standard native model. The increase in thermal stability results from the thermodynamic destabilization of the unfolded state (and consequent stabilization of the native one) due to entropy loss resulting from a restricted conformational space. It is also interesting to note that the *C_V_* curve for the WT form exhibits two peaks when the SS-bond is either stabilised (or locked), which is an indication that the protein no longer melts as a single unit, i.e., the melting transition is no longer two-state. It is important to mention that in line with experimental results, the thermal stability of ΔN6 is lower than that of the WT form. However, the model does not capture the decrease of thermal stability observed for the D76N mutant through *in vitro* experiments [12]. However, this is not relevant in the context of the present study, where we will compare the folding transition at the characteristic temperature (either *T_m_* or *T_f_*), of each model system, and within each model system.

**Figure 3.**
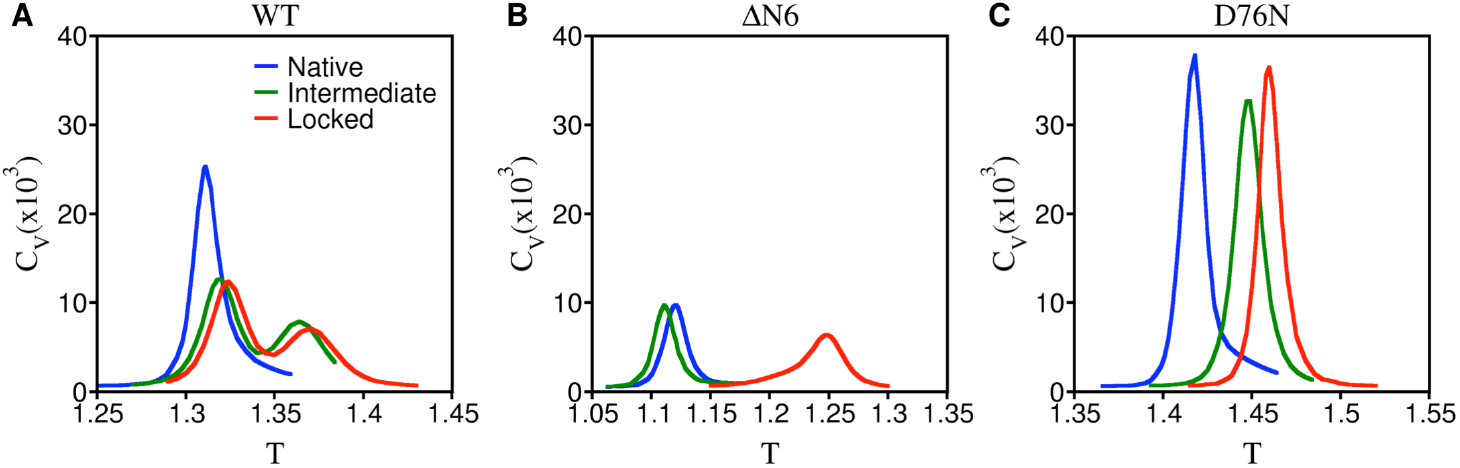
Thermal stability. Heat capacity as a function of *T* for WT form, ΔN6 structural variant, and D76N mutant. For each model system, three curves are shown corresponding to the three models of the SS-bond investigated in this study.

To explore the effect of the SS-bond on the folding transition, we evaluated the dependence of the energy on temperature. Specifically, we consider the energy normalised to the energy observed at *T_m_*, i.e., *E/E*(*T_m_*), and the temperature normalised to *T_m_* (Fig. 4 (A-C)). For the three model systems, the curve shows a clear sigmoidal shape when the SS-bond is modelled as a standard native interaction. However, the curve is clearly less steep for the ΔN6 mutant, which signals a less cooperative transition in this case. Locking the SS-bond produces no effect on the folding transition of the ΔN6 variant, has a minor effect on the folding transition of D76N mutant, but drastically changes the curve of the WT form, which is no longer sigmoidal when the SS-bond is locked. For the WT form, strengthening the SS-bond has thus a striking effect on the folding transition, which becomes clearly less cooperative. This smoothing of the folding transition foretells the formation of intermediates states across the folding pathway. Indeed, in previous studies in which the SS-bond was modelled as a standard native interaction, we found that the ΔN6 populates a folding intermediate state that we named intermediate *I* that preserves the nativelike core (extending from residue 21 to residue 94), conserves the trans isomerization of Pro32 characteristic of *I_T_*, and features an unstructured strand A that detaches significantly from the core [17]. In a posterior investigation [13], we found that both the WT form and D76N mutant, populate an intermediate state with an unstructured and detached C-terminus, that we named *I*_1_, while the D76N mutant exclusively populates another intermediate state, termed *I*_2_, with both termini unstructured and detached. Both *I*_1_ and *I*_2_ intermediates feature a nativelike core (*RMSD*_21-94_ < 2.0Å). In intermediate *I*_1_, 53% of the most hydrophobic residues are more solvent exposed than in the native structure, and this number increases to 62% in the intermediate *I*, and to 76% in *I*_2_. This enhancement in SASA for the hydrophobic residues implies that the identified states are potentially prone to aggregate, and their aggregation potential has been evaluated through proteinprotein docking simulations [14], with *I*_2_ standing out as a potentially important trigger of the aggregation cascade. The behaviour of the folding transition observed when the SS-bond is strengthened suggests that this bond alone may be enough to modulate the population of these intermediate states. In particular, are the previously observed intermediate states conserved, enhanced or suppressed when alternative models of the SS-bond are considered? Does a more stable SS-bond leads to the formation of novel intermediate states?

**Figure 4.**
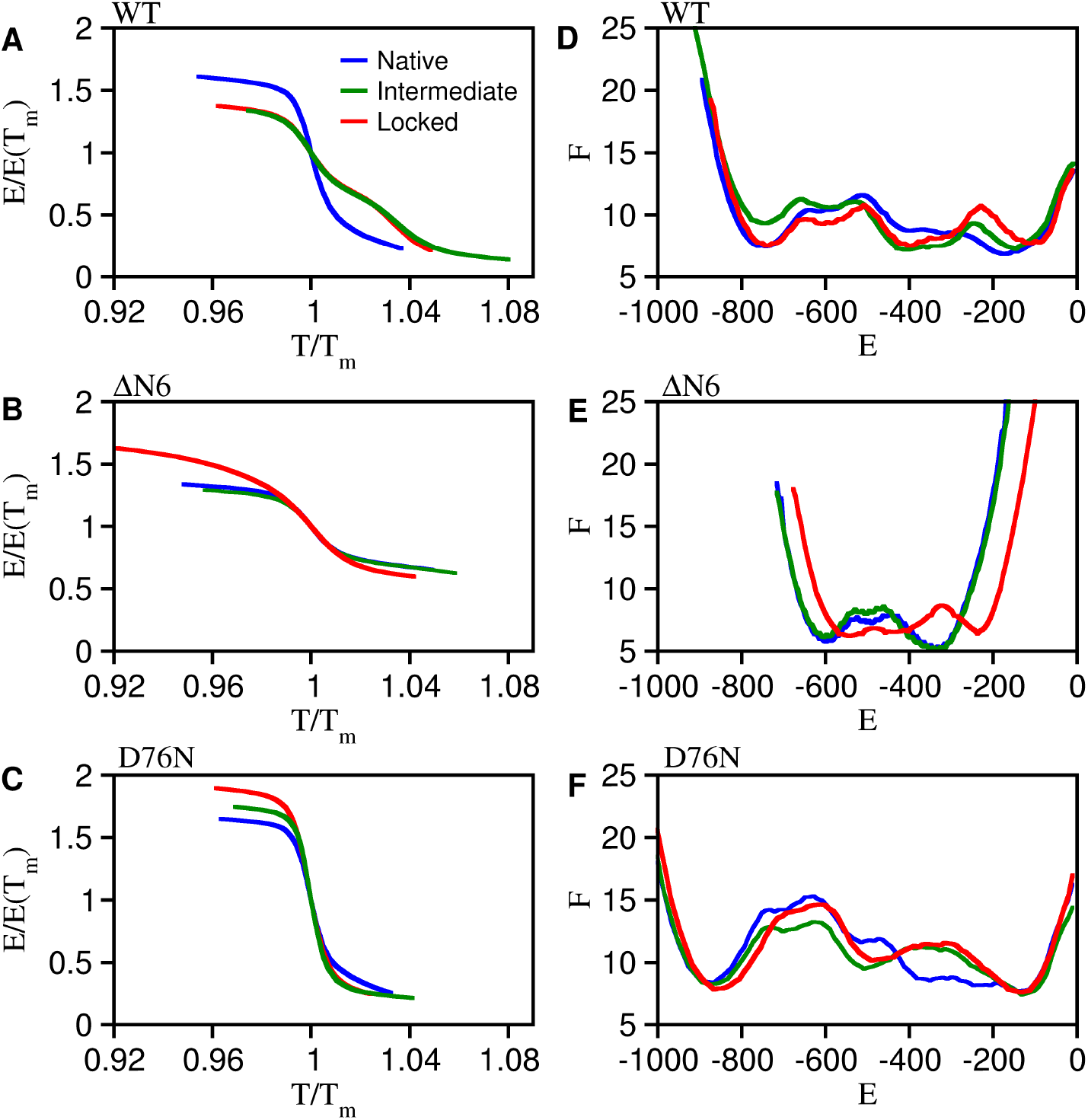
Folding transition. Temperature dependence of the energy normalised to the energy observed at *T_m_* for WT form (A), ΔN6 structural variant (B), and D76N mutant (C), and free energy projected on the energy *E* for the WT (D), ΔN6 (E), and D76N (F) at *T_f_*. For each model system, three curves are shown corresponding to the three models of the SS-bond investigated in this study.

### 3.3. Free energy profiles and surfaces

To explore the effect of the SS-bond on the population of intermediate states we start by analysing the free energy profiles, *F*(*E*), at the folding transition temperature, *T_f_* (Figure 4 (D-F)). The free energy profiles for the WT form are broadly conserved across the considered SS-bond models, but there is a novel intermediate state with energy *E* ≈ —400, which is compatible with intermediate *I*_2_ that was exclusively detected for the D76N mutant in our previous studies [13, 14]. In the case of the ΔN6 variant, the free energy profile corresponding to the locked interaction model is substantially different from the other two. There is a remarkably broad native basin encompassing the native state and, presumably, also the intermediate state *I*, and a considerable shift of the denatured basin towards higher energy. Since the locked SS-bond constraints the conformational space by blocking the formation of unfolded and extended conformations, the observed shift must result from the considerably higher temperatures used in the RE simulations of Δ*N*6 when the SS-bond is locked, which favour the population of high energy conformations. Finally, in the case of the D76N mutant, the basin corresponding to the intermediate state *I*_2_ (*E* ≈ —500) is deeper when the SS-bond is stabilised indicating a larger population (and, therefore, a higher thermodynamic stability) of this particular intermediate state.

To gain more information about the intermediate states, we looked into the free energy surfaces, i.e., the projection of the free energy on two reaction coordinates, the energy *E* and *R_g_* to the native structure (Figure 5), or the energy *E* and *RMSD* to the native structure (Figure 6). An immediately visible (and expected) effect of locking the SS-bond is the substantial shortening of the conformational space of the WT form and D76N mutant that no longer exhibits extended conformations (*R_g_* ≥ 35Å and *RMSD* ≥ 30Å). Interestingly, ΔN6 does not populate extended conformations, even when the SS-bond is modelled as a standard native bond. Indeed, the truncated variant exhibits a smaller conformational space in comparison with the other two model systems. This behaviour does not stem from poor sampling at higher temperatures since the *C_V_* tends to zero in the high temperature region above *T_m_* (Figure 3). Instead, it is intrinsically driven by the native contact map, that underlies protein folding energetics through the adopted Gō potential.

**Figure 5.**
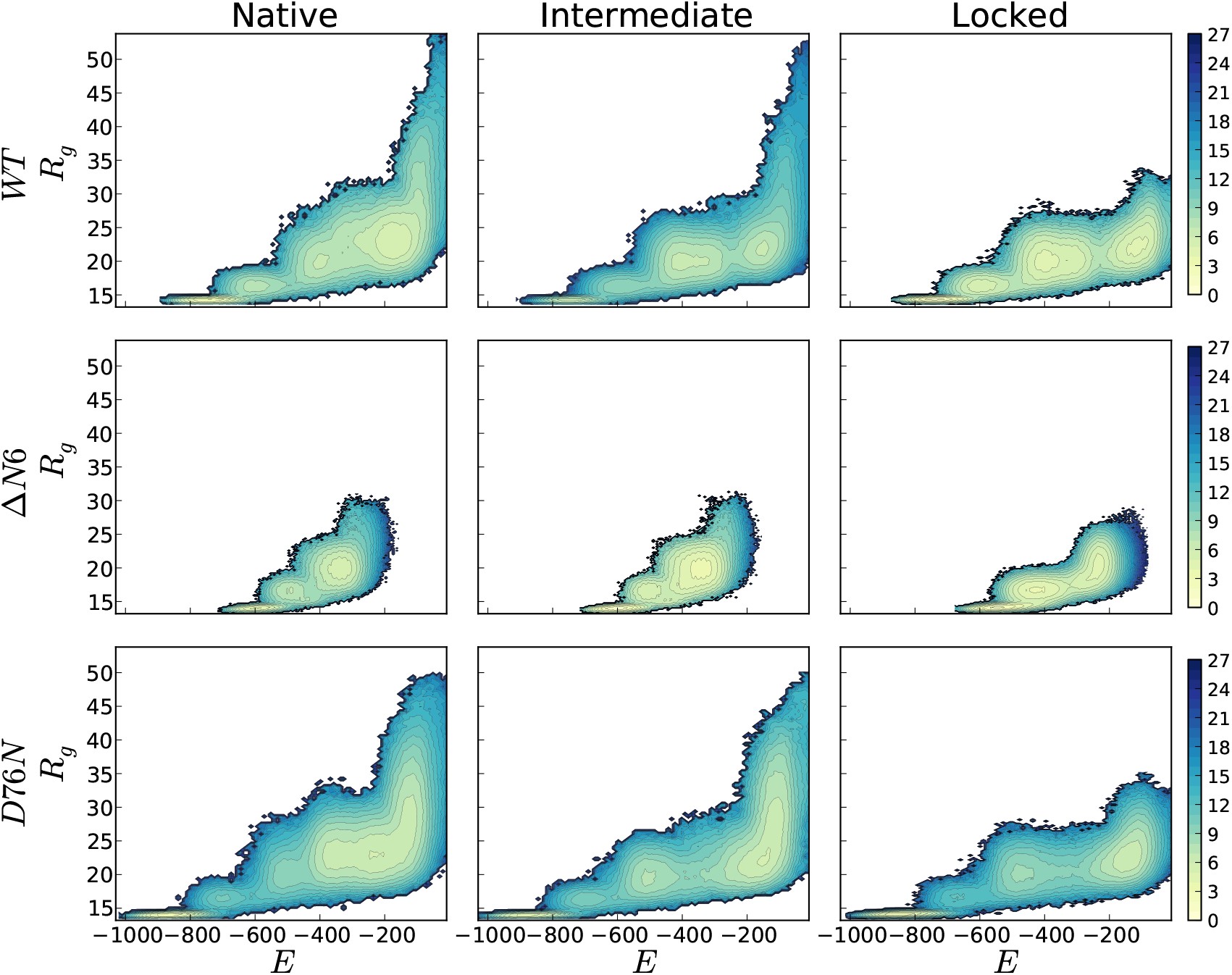
Free energy surfaces. Free energy projected on the energy *E* and *R_g_* for the WT form (top row), ΔN6 structural variant (middle row) and D76N (bottom row) at *T_f_*.

**Figure 6.**
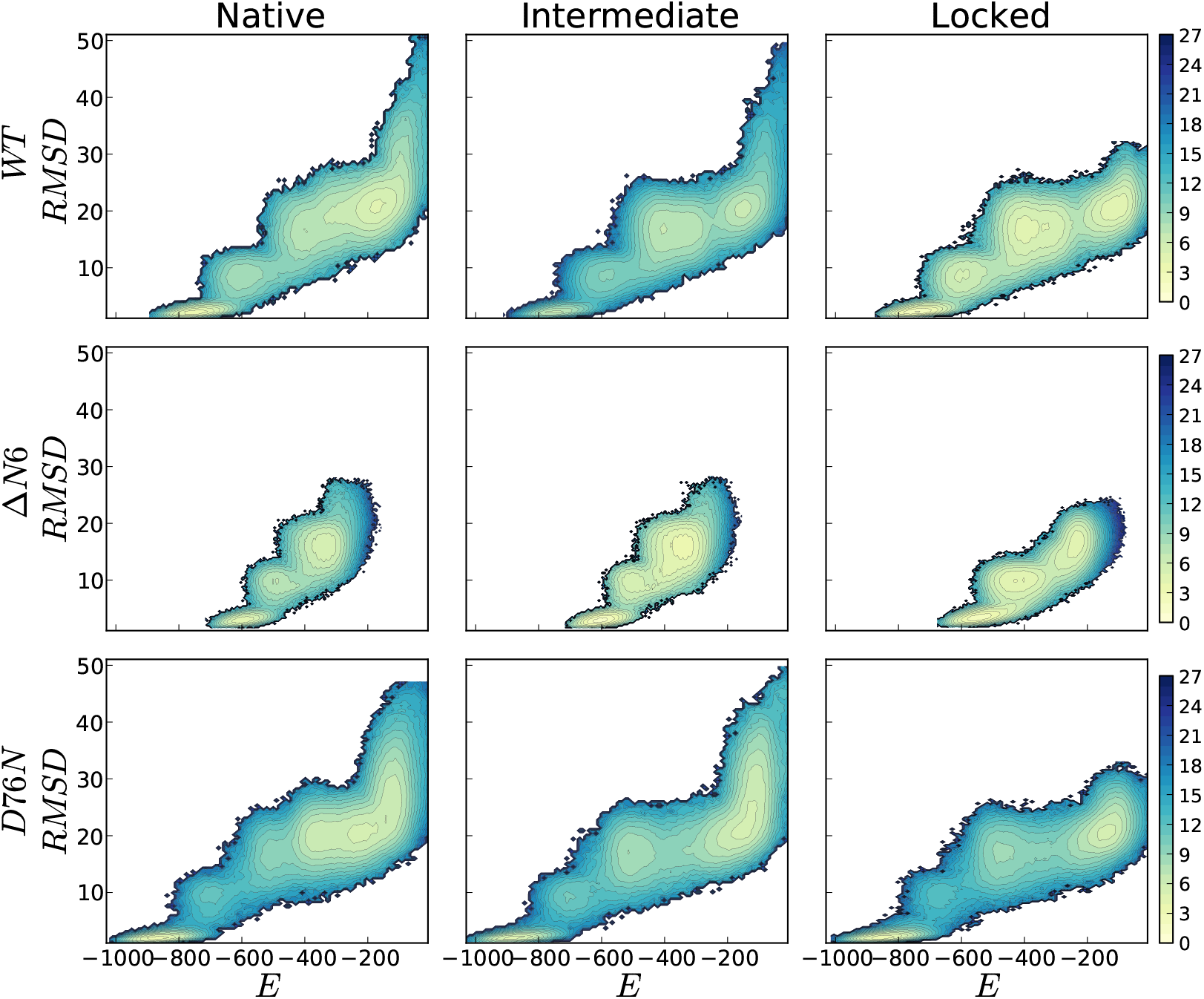
Free energy surfaces. Free energy projected on the energy *E* and *RMSD* for the WT form (top row), ΔN6 structural variant (middle row) and D76N (bottom row) at *T_f_*.

The analysis of the free energy surfaces indicate that irrespective of the considered model system a more stable, and eventually locked, SS-bond leads to considerably deeper free energy basins, and, therefore, to intermediate states with higher thermodynamic stability. In the case of the WT form, the analysis of the free energy surfaces confirms the population of an intermediate state (*E* —400; *R_g_* ≈ 20Å; *RMSD* ≈ 8Å) that is not present when the SS-bond is modelled as a standard native interaction. This intermediate state is also populated by the D76N mutant, and we anticipate that it should be *I*_2_, or some topological equivalent of *I*_2_. In line with what is observed in the free energy profiles, the intermediate basin in the free energy surface of the ΔN6 variant (*E* = —500; *R_g_* ≈ 17Å; *RMSD* ≈ 8Å) forms an almost continuum with the native basin, indicating that conformational transitions between the two states should be facilitated when the SS-bond is locked.

### 3.4. Identification of the intermediate states

Here we carry out an extensive clustering analysis in order to identify the intermediate states detected in the free energy surfaces of the three model systems. We recall that since the Gō potential is native centric, it is not able to accurately capture regions of the free energy landscape in which the conformations may be energetically stabilised by non-native interactions. We are particularly interested in determining if whether or not the WT form populates the intermediate with the two termini unstructured that was previously found for *D*76*N*. In order to do so, we performed structural clustering over the ensembles of conformations collected from the DMD simulations at fixed temperature. No novel intermediate states were detected for the ΔN6 and D76N mutant, but the clustering analysis reveals that that under the locked SS-bond model, the WT form indeed populates an intermediate state with the two termini unstructured and detached from the core that is identical to *I*_2_ and was not detected in previous studies based on the native bond model. In Figure 7 we report the representative conformation, together with the conformations corresponding to *I* and *I*_1_.

**Figure 7.**
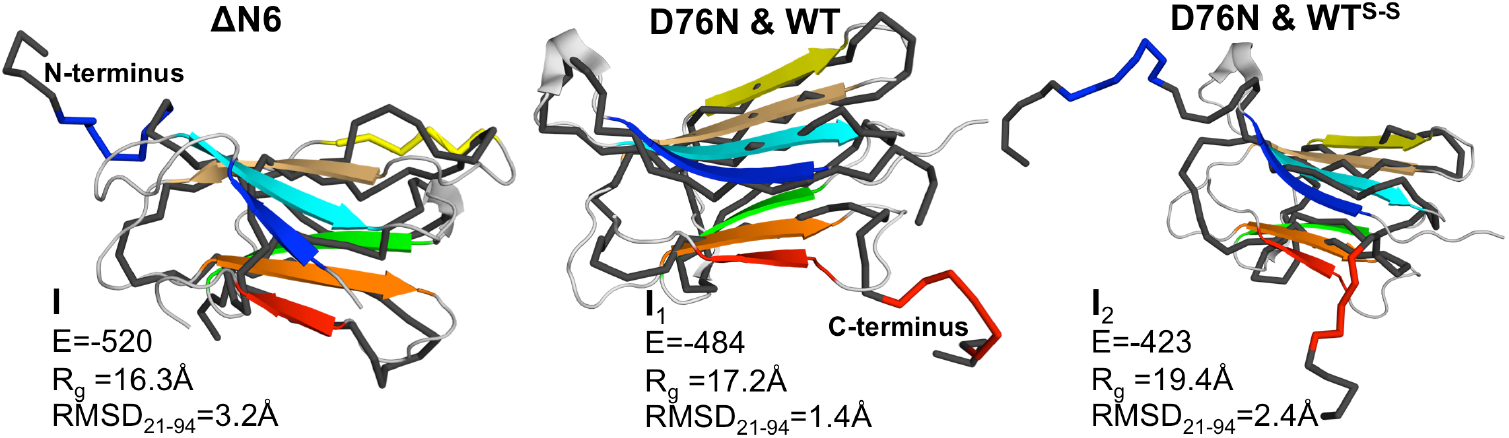
Intermediate states. Ribbon representation of the three-dimensional structure of the intermediate states *I*, *I*_1_ and *I*_2_ fitted onto the native structure of Δ*N*6 (I) and WT form (*I*_1_ and *I*_2_). In all the three cases the core (21 — 94) is well preserved (*RMSD* < 2.5Å). The intermediate I, featuring the N-terminus unstructured, is exclusively populated by the Δ*N*6 structural variant. The intermediate state *I*_1_, featuring the C-terminus unstructured, is populated by both the WT form and D76N mutant. The intermediate state *I*_2_, with both termini unstructured, is populated by the D76N mutant and by the WT form. However, in the latter case it only forms when the SS-bond is either stabilized relative to the other native interactions or locked. The intermediates *I*_1_ and I shown in this figure were found for the WT form when the SS-bond is locked.

## 4. Conclusions

Coarse-grained models with different levels of structural resolution have a long history and continue to be used in molecular simulation studies of protein folding [35]. Structurebased models such as the Gō model (reviewed in [36]) are particularly popular because they lead to smooth free energy landscapes, and encapsulate the importance of native structure as a driver of protein folding mechanisms and kinetics [37]. Since the Gō model is native-centric, it is natural to expect that simulated folding behaviour may be sensitive to the exact set of atomic coordinates that define the native structure. This question was originally addressed by Rey and co-workers in the context of a *C_α_* representation [38], [39], and later explored by one of us in the context of a full atomistic representation [19]. In the present work we explored the impact on folding of a particular aspect of the native structure of protein *β*2m, which is the existence of a SS-bond between residues 25 and 80. Evaluating the role of the SS-bond is particularly pertinent in the case of *β*2*m* since the protein has biomedical interest, and its aggregation behaviour appears to depend critically on the formation of the SS-bond. We considered three different model systems of *β*2m, namely, the WT form, the Δ*N*6 structural variant, and the D76N mutant, and found out that the SS-bond has an impact on the folding transition and thermodynamics of all of them, albeit to different extents. For the three model systems it is observed that a more stable, and eventually locked, SS-bond leads to a higher thermal stability of the native state, and also to a higher thermodynamic stabilization of the intermediates states that populate the folding pathway. Remarkably, a locked SS-bond has the most stringent effect on the folding transition of the WT form, which loses much of its cooperative character. Consistent with this behaviour, the WT form with the SS-bond locked populates an extra intermediate state featuring two unstructured termini and a well preserved core, which was not found in previous molecular simulation studies that modelled the SS-bond as a standard native interaction. Indeed, in that case only one intermediate with the C-terminus unstructured was detected [13]. The intermediate with two termini unstructured is also extensively populated by the D76N mutant [13], [40], and recent experimental studies indicate that it may play an important role as a trigger of the aggregation mechanism of D76N [18]. In fact it was found that the D76N mutant sparsely populates a highly dynamic conformation that exposes very aggregation prone regions as a result of a loss of *β*-structure at the N- and C-terminal strands, and the higher aggregation propensity of D76N was associated with the destabilization of its outer strands. So far, such an intermediate state was not observed *in vitro* for the WT form but one may not rule out the possibility that it may form *in vivo*. Indeed, it has been proposed that the presence of collagen in the osteoarticular tissues where *β*2m aggregates, creates not only a gradient of protein concentration that may favour self-association near the surface, but more importantly, it provides strong destabilizing electrostatic conditions that may trigger a spontaneous destructuring transition to a fibril-competent species, starting with the detachment of the N- and or C-terminal strands[41]. The results reported here, despite being merely predictive, are clearly in line with this hypothesis.

## Acknowledgments

The authors would like to thank Fundação para a Ciência e a Tecnologia (FCT), for financial support through grant number PTDC/FIS-OUT/28210/2017. This work was supported by the UIDB/04046/2020 Centre grants from FCT, Portugal (to BioISI).

## Author contributions

PFNF designed research, JM conducted the simulations, all authors analysed the data, PFNF and JM prepared the figures, PFNF wrote the article.

## Notes

### Competing Interest Statement

The authors have declared no competing interest.

